# *De novo* genome assemblies of threatened Asian hornbills (Bucerotidae) reveal declining population trajectories during the late Pleistocene

**DOI:** 10.1101/2025.05.20.655227

**Authors:** Pooja Yashwant Pawar, Gopi Krishnan, Rohit Naniwadekar, Jahnavi Joshi

## Abstract

Asian hornbills are flagship species of the wet tropics that face significant threats from hunting, habitat loss, and fragmentation. Despite being conservation flagships, whole genome information is available for only two of the 32 Asian hornbill species. In this study, we provide the first *de novo* genome assemblies for four hornbill species (Bucerotidae) in Asia. We present hybrid genome assemblies for *Buceros bicornis* and *Rhyticeros undulatus,* and short-read genomes for *Anthracoceros coronatus*, and *Aceros nipalensis*. The genome sizes of these hornbills range from 1.1 Gb to 1.3 Gb with over 93% completeness and gene prediction BUSCO. We also provide annotated mitogenomes for each of these species. Furthermore, we estimated Pleistocene population trajectories for these four species, which vary in their habitat preferences. Our results indicate that Pleistocene climatic fluctuations have led to dramatic population declines in all four species. We believe that this study provides robust genomic resources to support future conservation genomics efforts for hornbills.

## 3. INTRODUCTION

Hornbills (Order: Bucerotiformes) are avian flagships of tropical Asia and Africa, occupying a wide range of habitats ranging from African savannas to the wet evergreen forests of Southeast Asia^1^. While savanna-dwelling hornbills are primarily carnivorous, forest-dwelling species are predominantly frugivorous, playing a crucial role in seed dispersal and the regeneration of tropical forests^2^. Unfortunately, hornbills face severe threats from hunting and forest loss^3,4^, resulting in isolated populations that often occur in low densities^5–7^. Consequently, many Asian hornbill species, especially island endemics, are likely to have low genetic diversity due to small populations and isolation. For example, a recent mtDNA study on the point endemic Narcondam Hornbill revealed extremely low nucleotide diversity, highlighting the value of genomic tools in understanding hornbill populations^8^. While molecular phylogenetic studies have clarified evolutionary relationships among hornbills, some remain unresolved and could benefit from genomic work^9^. Genomic data could also provide information on molecular underpinning for trait variation in hornbills, which differ in morphology, colour, and vocalisations.

Partial mitogenome characterizations have been helpful for understanding genetic diversity and gene flow in endangered hornbills^10^. For example, mitogenomes revealed no genetic differentiation between captive Great Hornbills (*Buceros bicornis*) across five zoos in Thailand, highlighting their utility in ex-situ conservation^11^. Thus, genomic information can be critical for the management and conservation prioritisation of these rare species.

Interestingly, genomic data also allows for inferring the demographic histories of a species, which can provide valuable insights into how past climatic changes or anthropogenic disturbances might have influenced population dynamics. Rhinoceros Hornbill *Buceros rhinoceros* exhibited declines in effective population size during the Pleistocene^12^. Given that hornbills show strong habitat preferences and have been targeted by hunting for at least the last 50,000 years, they make an ideal candidate group for exploring population demography over historical time scales.

Of the 62 extant hornbill species, the genomes of the Rhinoceros Hornbill *Buceros rhinoceros* (Accession no. ASM71030v1), Great Hornbill (ASM2756380v1), and Ground Hornbill *Bucorvus abyssinicus* (ASM1339888v1) have been assembled using only short-read sequencing data. Additionally, the mitochondrial genomes of the Writhed-billed *Rhabdotorrhinus waldeni*, Visayan *Penelopides panini*, and Great and Wreathed Hornbills have been sequenced^13–15^.

Like other Asian Hornbills, Great Hornbill *Buceros bicornis* (hereafter GH), Wreathed Hornbill *Rhyticeros undulatus* (hereafter WH), Malabar Pied Hornbill *Anthracoceros coronatus* (hereafter MPH), and Rufous-necked Hornbill *Aceros nipalensis* (hereafter RNH) face alarming threats from rapid habitat loss, habitat destruction, hunting and illegal wildlife trade across their distribution ranges^2–4^. The GH, WH, and RNH are categorised as ‘Vulnerable’ in the IUCN Red List and are included in the CITES, while the MPH is classified as ‘Near Threatened’. Conserving these birds remains a challenge across their range-distribution countries. While the GH, WH, and RNH prefer evergreen forests, the MPH prefers the moist deciduous forests of the Indian subcontinent. Given their differences in habitat preferences, they offer contrasts for investigating population demographics in the historical context, especially since South and Southeast Asia have experienced significant fluctuations in the extent of wet and dry forests in the Pleistocene^16,17^.

In this study, we present assembled and annotated genomes of four Asian hornbill species: GH, WH, MPH, and RNH (Fig. 1). We used a combination of long-read and short-read sequencing data for GH and WH and short-read data for MPH and RNH. In addition to nuclear genomes, we annotated and compared their mitochondrial genomes. Finally, we used the Pairwise Sequentially Markovian Coalescent (PSMC) method^18^ to reconstruct their demographic histories and examine historical changes in the effective population size.

**Figure 1.**
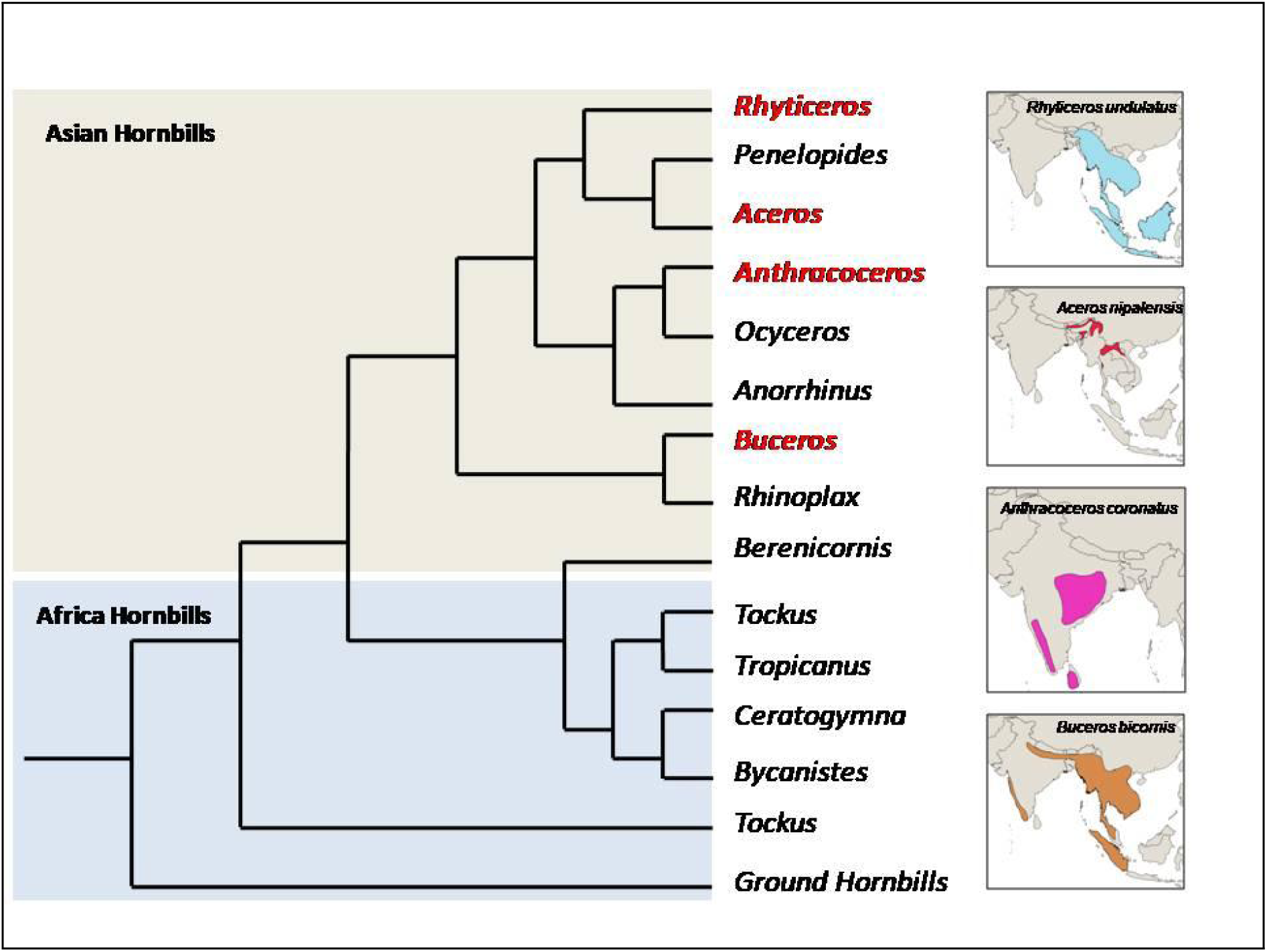
Thematic representation of phylogenetic relationships of hornbill genera (based on Gonzalez et., al. 2013) and geographic distribution ranges of four study species as per IUCN.

## 4. MATERIALS AND METHODS

### 4.1 Sample collection and DNA extraction

We collected tissue samples of GH, WH, RNH, and MPH. The GH sample was obtained from a deceased captive individual at Nehru Zoological Park, Hyderabad. One GH sample was from the Western Ghats in north Karnataka. WH samples were collected from trophies in eastern and western Arunachal Pradesh, India and the four RNH samples were collected from eastern, central and western Arunachal Pradesh in the Eastern Himalayan region.The MPH sample was collected from a deceased individual in the Western Ghats region of Maharashtra. Sample collection was carried out with prior approval from the Research Ethics Committee (NCF-EC-01/12/2022-(73)) and valid research permits from the state forest departments (CWL/GEN/2018-19/Pt.IX/NG/365-68; MSBB/Desk-5/Research/652/2023-24; Desk −22(8)/WL/CR-68(19-20)/397723-24, PCCF(WL)/E2lCR-1 1, 12022-23). We carried out high molecular weight DNA extraction using the Phenol-Chloroform-Isoamyl alcohol method for Oxford Nanopore sequencing (Appendix) and the QIAGEN DNeasy Blood & Tissue kit for Illumina sequencing.

### 4.2 Library preparation, sequencing, and read quality check

Library preparation for both Illumina and Oxford Nanopore platforms and sequencing was done at the Next Generation Sequencing (NGS) Facility at the CSIR-Centre for Cellular and Molecular Biology, Hyderabad, India. We generated long-read sequence data using the Oxford Nanopore (ONT) sequencing platform and short-read (150 bp paired-end) sequences using the Illumina NovaSeq 6000. We examined the sequencing quality for long-read and short-read data. The Phred quality score threshold was set at nine for long-read data. For the paired-end short-read data, we filtered out duplicates and low quality reads (sequences <50 bp, reads with gaps) using an accuracy threshold of Q20 with Fastp (v 0.23.4)^19^.

### 4.3 Genome assembly

We assembled the genome of GH and WH using long-read sequence data in Canu v. 2.3^20^. The genome assemblies obtained from ONT data were polished using corrected short-read data with Polca (from Masurca v. 4.1.0)^19,21^. We evaluated genome assembly completeness using Benchmarking Universal Single-Copy Orthologs (BUSCO v 5.7.1)^22^ with the Aves ortholog database (aves_odb10), which contains 8338 genes. The assembly statistics were generated using BlobToolKit (v 4.4.5)^23^, and the output was visualized as snail plots. For MPH and RNH, we assembled the genome using short-read data only. The paired-end short-read data was quality trimmed with FastP (v 0.23.2)^19^ and then assembled using Assembly By Short Sequences (ABySS v. 2.3.8), a de novo, parallel, paired-end short-read sequence assembler^24^. Similar to GH and WH, RNH genome assembly statistics were evaluated using BUSCO v 5.7.1 and BlobToolKit v 4.4.5^22,23^. We estimated the genome size using the overlap-layout-consensus (OLC) approach in Canu v. 2.3^20^ and the de Bruijn graph approach in ABySS v. 2.3.8^24^.

### 4.4 Repeat masking and genome annotation

To mask the repeat content of the genome, we performed *de novo* repeat prediction using RepeatModeler (v. 2.0.5)^25^. We built custom libraries for both genomes using the curated Dfam database and LTRStruct pipeline^25,26^. We then soft-masked the polished genomes with these custom libraries using RepeatMasker (v. 4.1.5)^26^.

Gene annotation was performed using GALBA (v. 1.0.11)^27^, which uses miniprot in combination with AUGUSTUS. For this, we created a reference protein dataset of 7,197,644 protein sequences of Aves from the Uniprot and Orthodb databases, which was used in the GALBA pipeline to predict protein-coding sequences in the soft-masked genome. Next, we removed unsupported predictions. We further filtered out multiple isoforms, retaining only the longest isoform for each gene, forming the final annotated gene dataset.

We evaluated the completeness of the gene annotation using BUSCO Orthologs (v 5.7.1) with the Aves ortholog database (aves_odb10), which contains 8,338 genes^22^. We further assessed the completeness and consistency of the gene annotation using OMArk (v. 2.0.3), with the whole OMA database (LUCA.h5) as reference^28^.

### 4.5 Single nucleotide polymorphisms and structural variants Heterozygosity

We compared the published *Buceros bicornis* genome from NCBI (ASM2756380v1) with the three genomes assembled in this study (*Buceros bicornis*, *Aceros nipalensis,* and *Rhyticeros undulatus*) to calculate single nucleotide polymorphisms (SNPs) and structural variants (SVs). Using default settings in MUMmer (v 3.23)^29^, we estimated SNPs and SVs by aligning each assembled genome with the ASM2756380v1 using Assemblytics (v1.2.1)^30^. Using BCFtools (v 1.21)^31^ and VCFtools (v 3.0)^32^, we calculated SNPs for 3 WH and 4 RNH individuals by mapping short-read data to the reference genomes assembled in this study.

### 4.6 Mitogenome assembly

We assembled and analyzed the mitogenome from short-read data with the packages GetOrganelle^33^, MITOS2^34^ web server and MitoZ (v. 3.6)^35^ to assemble, annotate and visualize mitogenomes of the four hornbill species. We estimated and compared GC content and GC skewness (G - C)/(G + C) across the total genome length, protein-coding genes, rRNA, and tRNAs for three mitogenomes assembled in this study and two published mitogenomes for *Buceros bicornis*^14^ and *Rhyticeros undulatus*^15^ to assess structural variation in the mitochondrial genome within and across species^36^.

### 4.7 Demographic history

We reconstructed demographic histories for all four species of hornbill using the pairwise sequentially Markovian coalescent (PSMC) method^18^. We removed scaffolds that mapped to the sex chromosomes from the genome assembly by comparing/referencing with the *Gallus gallus* genome. Illumina short-reads for 2 GH, 3 WH, 4 RNH and 1 MPH individuals were mapped to the autosome-only assembly of respective hornbill species, and variable sites were identified using BCFtools v 1.21^31^. We assessed the quality of mapping by estimating average genome coverage using QualiMap v 2.2.2^37^ The parameters for PSMC analysis were -N25 -t5 -r5 -p ‘25*2+4+4+6, where N is the number of iterations, t is the upper limit of time to the most recent common ancestor, r is the ratio of the scaled mutation rate and the recombination rate, and p is the number of free atomic time intervals. We bootstrapped the PSMC run 100 times to estimate the variance in effective population sizes. Generation time used for GH, WH, MPH, and RNH was 14.1, 8.6, 5.3, and 9.5 years^38^, respectively; and the mutation rate was 7×10^−9^ mutations per generation^12^. We performed PSMC analysis on multiple individuals of GH (n=2), WH (n=3) and RNH (n=4). One GH individual from the Western Ghats and another from the Eastern Himalaya were sampled. Two WH individuals from eastern and western Arunachal Pradesh each and four RNH individuals were sampled from eastern, western and central Arunachal Pradesh in the Eastern Himalayan region. The MPH was sampled from the Western Ghats. See supplementary materials for details of software, versions, and commands for genome assembly, and annotation.

## 5. RESULTS

### 5.1 Genome Sequencing

For GH, we obtained 26x coverage of long-read data with 4 million reads. A total of 436 million paired-end short reads were generated. After removing 9.3% duplications, we retained 429 million reads, resulting in 63x coverage of the short-read data. The mean read length was 148 bp, and the GC content was 45.49%, with 96.3% of bases having a Phred quality score greater than or equal to Q20. Similarly, for WH, MPH, and RNH, we obtained 65x, 119x, and 66.6x coverage of short-read data, respectively. The mean read length was 148 bp, and the number of reads generated was 442 million for WH, 882 million for MPH, and 493 million for RNH. The GC content across all three species was 45%. The long reads obtained for WH were 9.5 million, resulting in 54x coverage.

### 5.2 Whole genome assembly

The genome size of the *de novo* assembled GH, WH, MPH, and RNH genomes were 1.14 Gb, 1.16 Gb, 1.3 Gb, and 1.1 Gb, respectively (Fig. 2). The total number of assembled contigs was 2,192 for GH and 3,114 for WH, with contig N50 values being 6.3 Mb for GH and 11.3 Mb for WH, suggesting that both assemblies are highly contiguous. BUSCO evaluation to assess the genome completeness predicted the polished assembly of GH to be 94.8% complete and WH assembly to be 99.1% complete. The detailed assembly statistics are given in Table 1. In contrast, the MPH and RNH short-read-only genome assemblies were fragmented and had 1,162,667, and 422,266 scaffolds with scaffold N50 values of 231.9 Kb and 7.7 Kb, respectively. The BUSCO completeness score of the RNH genome was 93.2%. We could not obtain the BUSCO evaluation for MPH due to the high number of contigs in the genome assembly.

**Figure 2.**
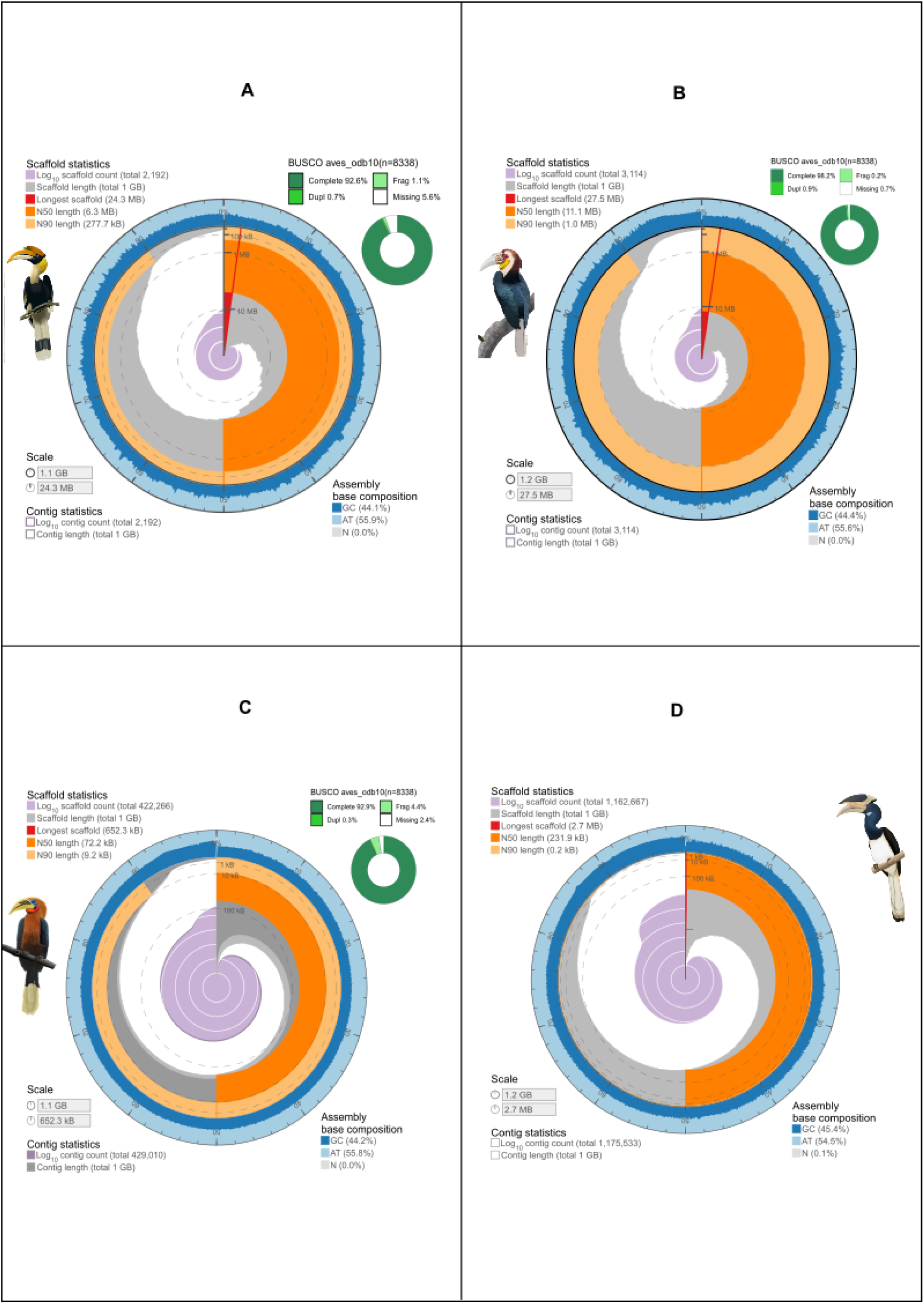
Genome assembly snailplots showing assembly stats sequence composition and BUSCO completeness score for A) Great Hornbill, B) Wreathed Hornbill, C) Rufous-necked Hornbill, and D) Malabar Pied Hornbill, respectively. (Hornbill illustrations by Bhagyashri Patwardhan)

**Table 1.** Genome assembly statistics for four species of hornbills.

|  | Great<br>Hornbill<br>(GH) | Wreathed<br>Hornbill<br>(WH) | Rufous-necked<br>Hornbill<br>(RNH) | Malabar Pied<br>Hornbill<br>(MPH) |
| --- | --- | --- | --- | --- |
| ***Genome assembly statistic*** |  |  |  |  |
| No. of long reads | 4 M | 9.5 M | - | - |
| No. of short reads | 429 M | 442 M | 493 M | 882 M |
| Coverage | 22.5 x | 34.21 x | 66.6 x | 199 x |
| Genome size | 1.14 Gb | 1.16 Gb | 1.1 Gb | 1.3 Gb |
| Number of contigs | 2192 | 3114 | 422266 | 1162667 |
| N50 | 6.3 Mb | 11.3 Mb | 7.7 Kb | 231.9 Kb |
| Largest contig length | 24.3 Mb | 27.5 Mb | 0.65 Mb | 2.7 Mb |
| GC content | 44.08 % | 44.41 % | 44.2 % | 45.4 % |
| % single copy genes<br>identified using BUSCO | 92.6 % | 94.9 % | 92.9 % | - |
| BUSCO completeness | 94.8 % | 99.1 % | 93.2 % | - |
| ***Repeat masking statistics (%)*** |  |  |  |  |
| Retroelements | 6.05 | 7.83 | 4.87 | 4.92 |
| -SINEs | 0.04 | 0.05 | 0.03 | 0.07 |
| -LINEs | 4.03 | 3.83 | 3.68 | 3.63 |
| DNA transposons | 0.10 | 0.07 | 0.13 | 0.10 |
| Unclassified | 2.89 | 2.03 | 4.09 | 10.79 |
| Small RNA | 0.37 | 0.15 | 0.02 | 0.05 |
| Satellites | 0.03 | 0.04 | 0.03 | 0.05 |
| **Total bases masked** | **9.54** | **10.13** | **9.12** | **15.88** |

### 5.3 Annotation

In repeat masking, 9.54%, 10.13%, 15.88% and 9.12% of genome sequences were masked for GH, WH, MPH, and RNH, respectively. The repeat regions were predominantly composed of Retroelements, accounting for 6.05 %, 7.83%, 4.92%, and 4.87% of the genomes in GH, WH, and RNH, respectively. The composition of repeat content and the detailed repeat statistics are given in Table 1.

GALBA predicted 21,316 and 21,357 protein-coding genes for GH and WH, respectively. BUSCO analysis to assess the completeness of protein predictions showed that the annotations were 93.3% complete for GH and 95.9% for WH. Similarly, the OMArk-based completeness assessment using 11,063 conserved Hierarchical Orthologous Groups estimated annotation completeness to be 94.63% for GH and 94.6% for WH. See Supplementary Table 7 for detailed OMArk statistics.

### 5.4 Single nucleotide variation and structural variants

We identified single nucleotide variations (SNPs) and structural variants (SVs) across all the hornbill genomes by comparing them against the published *B. bicornis* reference genome (ASM2756380v1) (Supplementary Table 1). In GH, we identified over 1.9 million SNPs and 22,896 SVs comprising 15.6 Mbp of the genome. The predominant structural variation was repeat contractions (50-500 bp), with a total count of 4,938. By comparing with the published *B. bicornis* (ASM2756380v1) genome, for WH, we identified over 53 million SNPs and 49,983 SVs, amounting to 38.4 Mbp, for RNH, we identified 52 million SNPs and 44,103 SV, and for MPH we identified 63 million SNPs and 30,778 SVs amounting to 5.5 Mbp. In the genomes of these four species, deletions (50-500 bp) were the most common type of structural variation, with total counts ranging between 11,213 to 12,724. On mapping individual data to the species-specific reference genomes, we obtained 1.7 million SNPs for RNH (n = 4), 2.8 million SNPs for GH (n = 2) and 10 million SNPs for WH individuals (n = 3).

### 5.5 Mitogenome assembly

The mitochondrial genome assemblies for GH, WH, MPH, and RNH were 15,552 bp, 15,794 bp, 15,569 bp, and 15,622 bp in size, respectively. All four mitogenomes comprised 13 protein-coding genes, 2 rRNAs, and 22 tRNAs (Supplementary Fig. 1). For the detailed mitogenome organization see supplementary Table 2-6. The overall GC and A + T content for the GH mitogenome were 46.9% and 53.1%, respectively. For WH, the GC and A+T content was 48.7% and 51.3%, respectively, for MPH, 47.7% and 52.3%, and for RNH, 48.6% of GC and 51.4%, respectively. The GC content was similar to the concatenated composition of the protein-coding genes, tRNA, and rRNA for the respective hornbill species. The mitogenomic structures were highly conserved between and across species as we did not see differences in GC contents of protein-coding genes, rRNA, tRNA and their respective GC skewness (Table 2.).

**Table 2.**
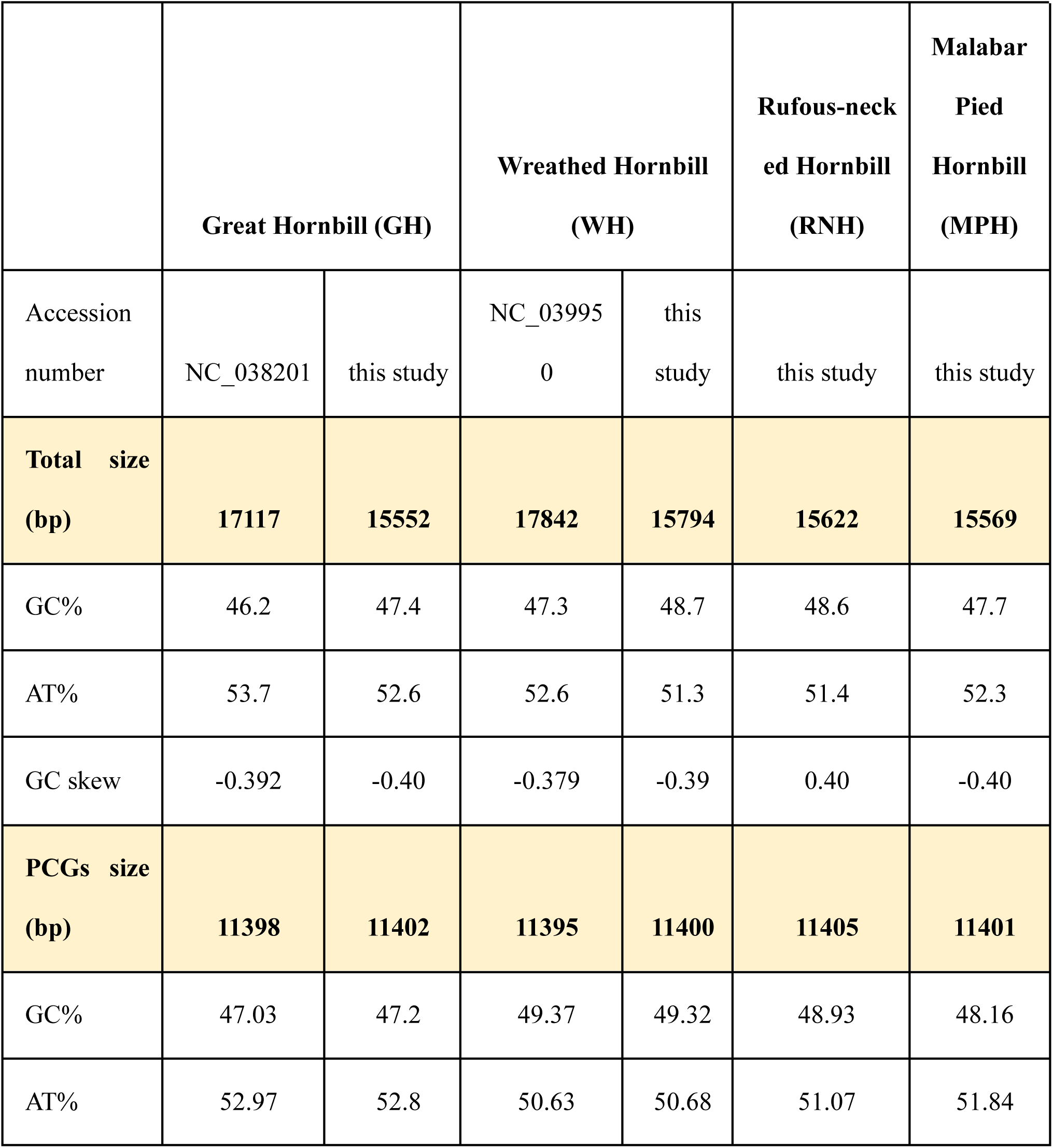
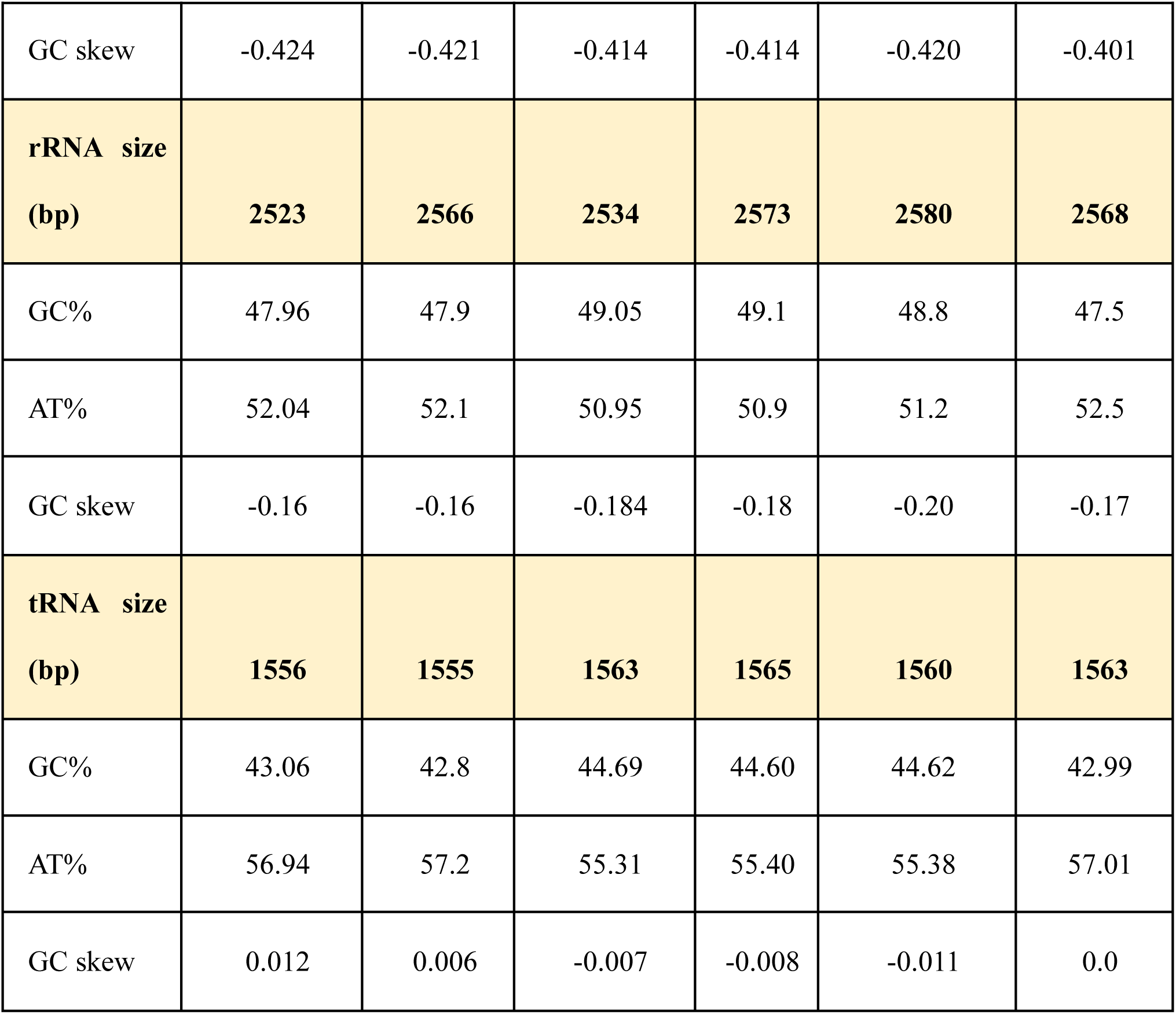
Intra- and inter-specific comparison of mitogenomic structures in four hornbill species.

|  | **Great Hornbill (GH)** |  | **Wreathed Hornbill (WH)** |  | **Rufous-neck ed Hornbill (RNH)** | **Malabar Pied Hornbill (MPH)** |
| --- | --- | --- | --- | --- | --- | --- |
| Accession number | NC_038201 | this study | NC_039950 | this study | this study | this study |
| **Total size (bp)** | **17117** | **15552** | **17842** | **15794** | **15622** | **15569** |
| GC% | 46.2 | 47.4 | 47.3 | 48.7 | 48.6 | 47.7 |
| AT% | 53.7 | 52.6 | 52.6 | 51.3 | 51.4 | 52.3 |
| GC skew | -0.392 | -0.40 | -0.379 | -0.39 | 0.40 | -0.40 |
| **PCGs size (bp)** | **11398** | **11402** | **11395** | **11400** | **11405** | **11401** |
| GC% | 47.03 | 47.2 | 49.37 | 49.32 | 48.93 | 48.16 |
| AT% | 52.97 | 52.8 | 50.63 | 50.68 | 51.07 | 51.84 |
| GC skew | -0.424 | -0.421 | -0.414 | -0.414 | -0.420 | -0.401 |
| **rRNA size**<br>**(bp)** | **2523** | **2566** | **2534** | **2573** | **2580** | **2568** |
| GC% | 47.96 | 47.9 | 49.05 | 49.1 | 48.8 | 47.5 |
| AT% | 52.04 | 52.1 | 50.95 | 50.9 | 51.2 | 52.5 |
| GC skew | -0.16 | -0.16 | -0.184 | -0.18 | -0.20 | -0.17 |
| **tRNA size**<br>**(bp)** | **1556** | **1555** | **1563** | **1565** | **1560** | **1563** |
| GC% | 43.06 | 42.8 | 44.69 | 44.60 | 44.62 | 42.99 |
| AT% | 56.94 | 57.2 | 55.31 | 55.40 | 55.38 | 57.01 |
| GC skew | 0.012 | 0.006 | -0.007 | -0.008 | -0.011 | 0.0 |

### 5.6 Demographic history

We estimated the demographic histories of GH, WH, MPH, and RNH using genome assemblies We observed notable differences in the demographic histories of four hornbill species (Fig. 3).

**Figure 3.**
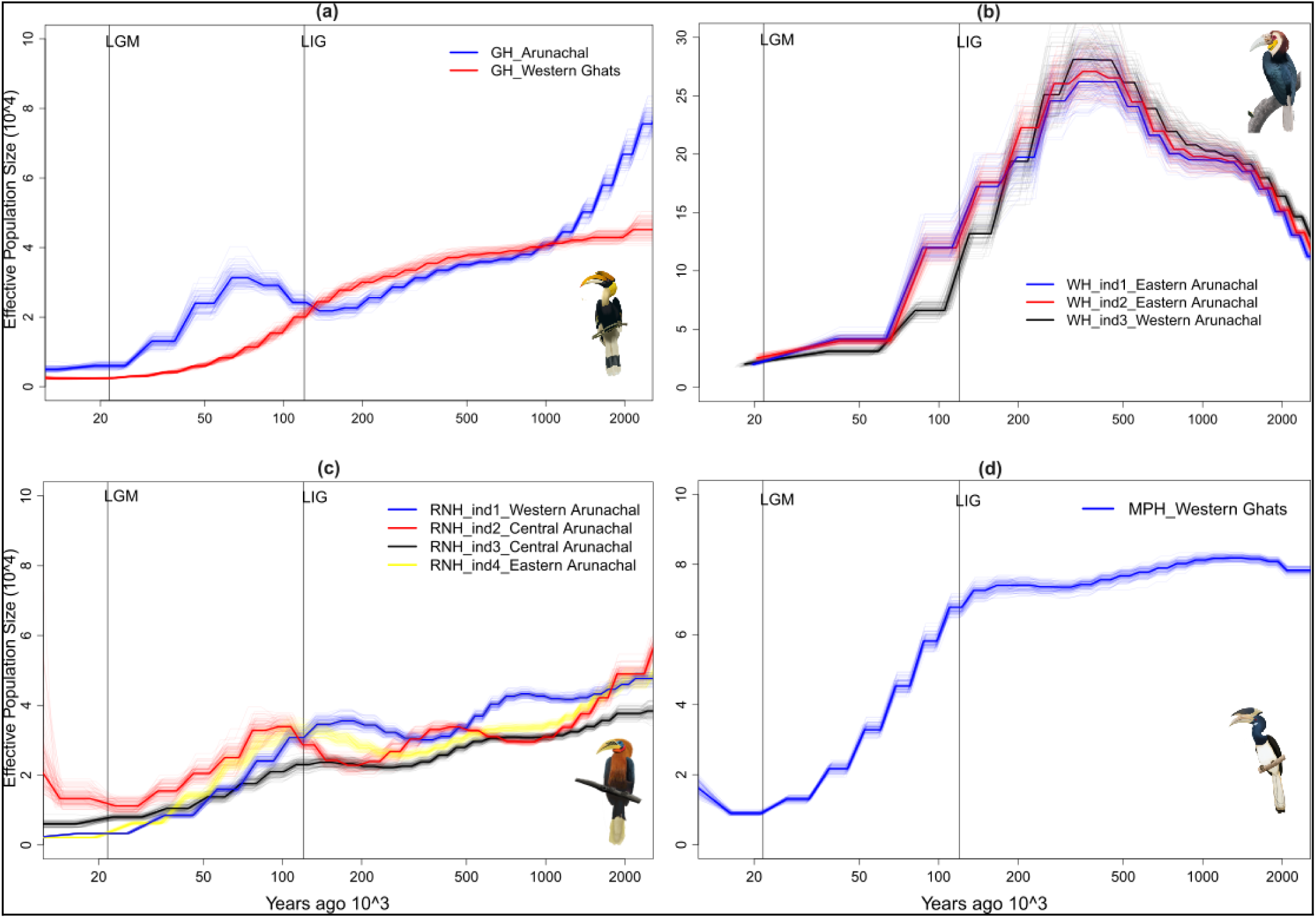
Demographic trends in effective population sizes (Ne) for four hornbill species. The grey box indicates the Pleistocene epoch (2.5 mya to 11.7 kya), and the vertical lines indicate the beginning of the last interglacial (LIG) and last glacial maximum (LGM). a) Great Hornbill *Buceros bicornis*; b) Wreathed Hornbill *Rhyticeros undulatus*; c) Rufous-necked Hornbill *Aceros nipalensis*; and d) Malabar Pied Hornbill *Anthracoceros coronatus.* (Hornbill illustrations by Bhagyashri Patwardhan).

For GH, the effective population size (Ne) estimates around 15 kya were 12,782 individuals in the Himalayan population and 7,755 individuals in the Western Ghats population. The GH populations in the Himalaya and the Western Ghats reached their peak Ne at the beginning of the Pleistocene ((2.5 mya), with 89,138 and 45,157 individuals, respectively (Fig 3a). For WH, Ne estimates in eastern Arunachal Pradesh ranged between 31,936 and 36,252 individuals at around 20 kya, while the western Arunachal population had a Ne of 36,049 individuals. The effective population sizes for WH peaked at ∼ 0.4 mya, with populations ranging from 2,62,083 to 2,81,166 individuals (Fig. 3b). It was followed by sharp declines for all three individuals throughout the last glacial period (120 kya to 11.9 kya). The RNH populations in eastern, central and western Arunachal Pradesh exhibited similar population trajectories, with peak Ne values between 38,422 and 60,450 individuals occurring approximately 2.5 million years ago (Fig. 3c). The MPH population from the Western Ghats had a Ne of 22,591 individuals at around 15 kya. The MPH population exhibited a relatively stable Ne trajectory until the Last Interglacial (LIG, ∼120 kya), peaking with 81,833 individuals.

## 6. DISCUSSION

The high-quality, annotated genome assemblies of the four hornbill species provided here will be valuable for future evolutionary and population genomics research on hornbills throughout Asia and Africa. The genome assemblies are robust with moderate to high sequencing coverages and exhibit high completeness. These are among the first hybrid genome assemblies of GH and WH. The contig N50 values or GH and WH exceed the average scaffold N50 reported for published bird genomes^39^. The total genome lengths for all four species fell well within the expected genome size (∼1-2.1 Gb) for birds^39,40^. The genome lengths of Asian hornbills presented in this study are comparable to the genomes of hornbill species (1.1 Gb) in Africa^41^ and Asia (NCBI: ASM71030v1, ASM2756380v1).

The newly assembled mitogenomes are circular and similar in size to those previously reported^14,15^. The gene lengths, and mitogenome organisations for GH and WH were highly comparable with the published mitogenomes^14,15^. When compared across species, the gene arrangements are highly conserved. However, the gene lengths showed slight variation in a few genes. This intra-species and inter-species mitogenome comparison provides a basis for insights into gene variation, showing evidence of mitochondrial structure diversity.

The relative prevalence of transpose elements (retroelements and DNA transposons) in all four hornbill genomes was similar to that of other bird species^39^. The mitogenome assembly sizes in hornbills from this study are comparable to the average for birds (mean = 15,464; range: 13,000-17,000; n = 363 bird species) and for Bucerotiformes (13,487-16,045)^39^. The BUSCO scores for gene predictions using GALBA were high (over 94%), indicating a high level of completeness. Additionally, OMARK placed the three genomes in this study under the Neognathae lineage, suggesting that annotated genomes are robust and reliable. Unfortunately, we were unable to verify this for the MPH, as the genome was fragmented.

We observed notable differences in the demographic histories of the four hornbill species. The peak effective population sizes differed across species over the analysed time span, with the WH having at least three times higher peak effective population sizes than its other Himalayan counterparts, the GH and RNH. A previous study found no phylogenetic signal in effective population sizes^12^, suggesting that closely related species may differ in their effective population sizes.

The different hornbill species responded idiosyncratically to multiple cooling cycles during the Pleistocene. During the mid-Pleistocene (till 400 kya), the Eastern Himalayan region likely maintained humid tropical conditions^42^, providing suitable breeding habitats for the WH and supporting a drastic increase in its effective population size. In contrast, GH in the Eastern Himalaya region showed a decline in population size during this time despite similar ecological preferences, highlighting idiosyncratic species responses to the environment.

Glacial periods, particularly during the onset of LGM (∼120 kya), were associated with reduced wet tropical forest cover, constraining habitat availability for these forest-dwelling birds. Populations of the MPH and GH in the Western Ghats and RNH in the Eastern Himalaya showed gradual declines, a trend observed globally for multiple bird species^12^.

Interestingly, the drastic declines in the Eastern Himalayan GH population (∼70 kya) aligns with regional evidence for a cool and dry climate^43,44^; however, an earlier sharp decline in the WH around 400 kya may reflect undocumented vegetation shifts. The RNH and, to a lesser extent, the MPH and GH from the Eastern Himalaya exhibited multiple demographic fluctuations, some pre-dating the LIC, pointing to complex interactions between climate, habitat and species-specific histories.

The Late Pleistocene vertebrate assemblages confirm the consistent occurrences of birds, including raptors and hornbills, from 126 kya to 11.7 kya in Southeast Asia^45,46^. Hornbill bones were found in the Niah cave excavations, dating back to 45 ka^47^. Hornbills also have traits such as large body size, communal roosting, and incarcerated nesting, making them vulnerable to hunting that resulted in extinction of some bird species^48^. Even relatively low levels of hunting pressures can cause local extirpations of hornbills. How much this contributed to the reduction in effective population sizes in the late Pleistocene remains to be understood.

## 7. CONCLUSIONS

The reference genomes of the *Buceros bicornis*, *Rhyticeros undulatus*, *Anthracoceros coronatus*, and *Aceros nipalensis* will serve as a resource to further understand the genetic makeup of hornbills. Using these reference genomes, we could reconstruct the demographic histories of these threatened species and understand their population trajectories in response to drivers such as climatic fluctuations and human footprint. It now enables us to delve into deeper inquiries in comparative genomics and assess genomic variation in the light of a spatio-temporal framework. With the increasing application of genomics into conservation biology, we hope that these reference genomes will act as a tool for conservation prioritization exercises and management for hornbills in the near future.

## Supporting information

pawar_et_al_suppl_hornbill_genomes

## 8. ACKNOWLEDGEMENTS

We thank the Arunachal Pradesh, Telangana and Maharashtra State Forest Departments for permitting us to conduct this study. We are grateful to the NGS and High-Perform-Cluster facility at the CSIR-Center for Cellular and Molecular Biology, Hyderabad. The study was funded by a start-up grant to Jahnavi Joshi and a SERB grant (No: SRG/2021/001523) to Rohit Naniwadekar. Pooja Yashwant Pawar was supported by the DST-WOSA (WOS-A/LS-278/2021) and the Rainmatter Foundation Fellowship. Gopi Krishnan was supported by the CSIR-NET fellowship. We thank Subhadra Devi (IFS) and Dr B Laxmi Narayana from the Nehru Zoological Park, Hyderabad, Japang Pansa, Himanshu Lad, and Vishal Sadekar for their support during the fieldwork. We thank Pragyadeep Roy, Vinay K L, Bharti Dharapuram, Vinay Sagar, Meghana Natesh, V. V. Robin and Kritika Garg for the discussions. We thank Bhagyashri Patwardhan for the illustrations of the four hornbill species.

## Author contributions

Idea: JJ; Conceptualization: PYP, JJ, RN; Funding: JJ, RN; Fieldwork: PYP, RN; Wet lab work: PYP; Data analysis: PYP, GK; Manuscript writing: PYP, RN; Comments & reviewing manuscript: PYP, RN, GK, JJ

## Data availability

The genome assemblies and raw sequence data will be submitted to the National Centre for Biotechnology Information (NCBI) genome repository.

