## Supplementary material for "*De novo* genome assemblies of threatened Asian hornbills (Bucerotidae) reveal declining population trajectories during the late Pleistocene": pawar_et_al_suppl_hornbill_genomes

Title: *de novo* genome assemblies for threatened Asian hornbills (Bucerotidae) reveal population declines during Pleistocene

Authors: Pooja Yashwant Pawar[1,2], Gopi Krishnan[3,4], Rohit Naniwadekar[2] and Jahnavi Joshi[3,4]

1 Manipal Academy of Higher Education, Manipal, India

2 Nature Conservation Foundation, Mysore, India

3 CSIR- Centre for Cellular and Molecular Biology, Hyderabad, India

4 Academy of Scientific and Innovative Research (AcSIR), Ghaziabad, India

Supplementary materials

**Supplementary Section 1: DNA extraction protocol**

High Molecular Weight DNA extraction protocol

Reagents required:

| **Reagent** | **Concentration** |
|---|---|
| Tris- HCl (pH=8) | 50 mM |
| EDTA | 100 mM |
| NaCl | 5 M |
| SDS | 20% |
| Proteinase K | 20 mg/ml |
| TE | 1x |
| Phenol, Chloroform, Isoamyl alcohol, Isopropanol | 25:24:1 |

Procedure:

1. Cut 25-50 mg of tissue into small pieces

2. Add 1 volume of lysis buffer (50 mM Tris-HCl pH 8.0, 100 mM EDTA, 100 mM NaCl), 100 ul of 20% SDS, 20 ul of 20 mg/ml Proteinase K.

3. Lyse for overnight (optional) at 56°C with low RPM.

4. Transfer the supernatant to a new 2ml tube.

5. Add 1 volume of Phenol:Chloroform: Isoamyl alcohol (25:24:1 ratio). Incubate on a rotating wheel

for 10 mins.

6. Centrifuge at 10,000 RPM for 10 minutes. Transfer the aqueous (upper) phase to a new 2ml tube.

7. Repeat steps 5–6.

8. Add 1 volume of Chloroform: Isoamyl alcohol (24:1 ratio). Incubate on a rotating wheel for 10

mins.

9. Centrifuge at 10,000 RPM for 10 minutes. Transfer the aqueous (upper) phase to a new 2ml tube.

10. Add 50 ul of 5M NaCl and one volume of chilled isopropanol.

11. Keep overnight precipitation at -20°C.

12. Centrifuge at 10,000 RPM for 15 minutes. Remove supernatant.

13. Wash the pellet twice with 1 volume of 70% ethanol followed by centrifugation.

14. Dry spin the columns for 1 minute at 10,000 RPM.

15. Air-dry the pellet for 5-10 minutes at room temperature.

16. Dissolve the pellet in 1 x TE prewarmed to 55°C. Store DNA at 4°C.

**Supplementary Section 2: NCBI Assembly accession numbers (Will be updated once published)**

| Accession number | Species | Sex | Sample type | Assembly type |
|---|---|---|---|---|
| | *Buceros bicornis* | Female | tissue | Hybrid de novo |
| | *Rhyticeros undulatus* | unknown | trophy | Hybrid de novo |
| | *Aceros nipalensis* | unknown | trophy | Short-read only de novo |
| | *Anthracoceros coronatus* | unknown | tissue | Short-read only de novo |

**Supplementary Section 3: Genome assembly and annotation workflow with tools used shown in green boxes**

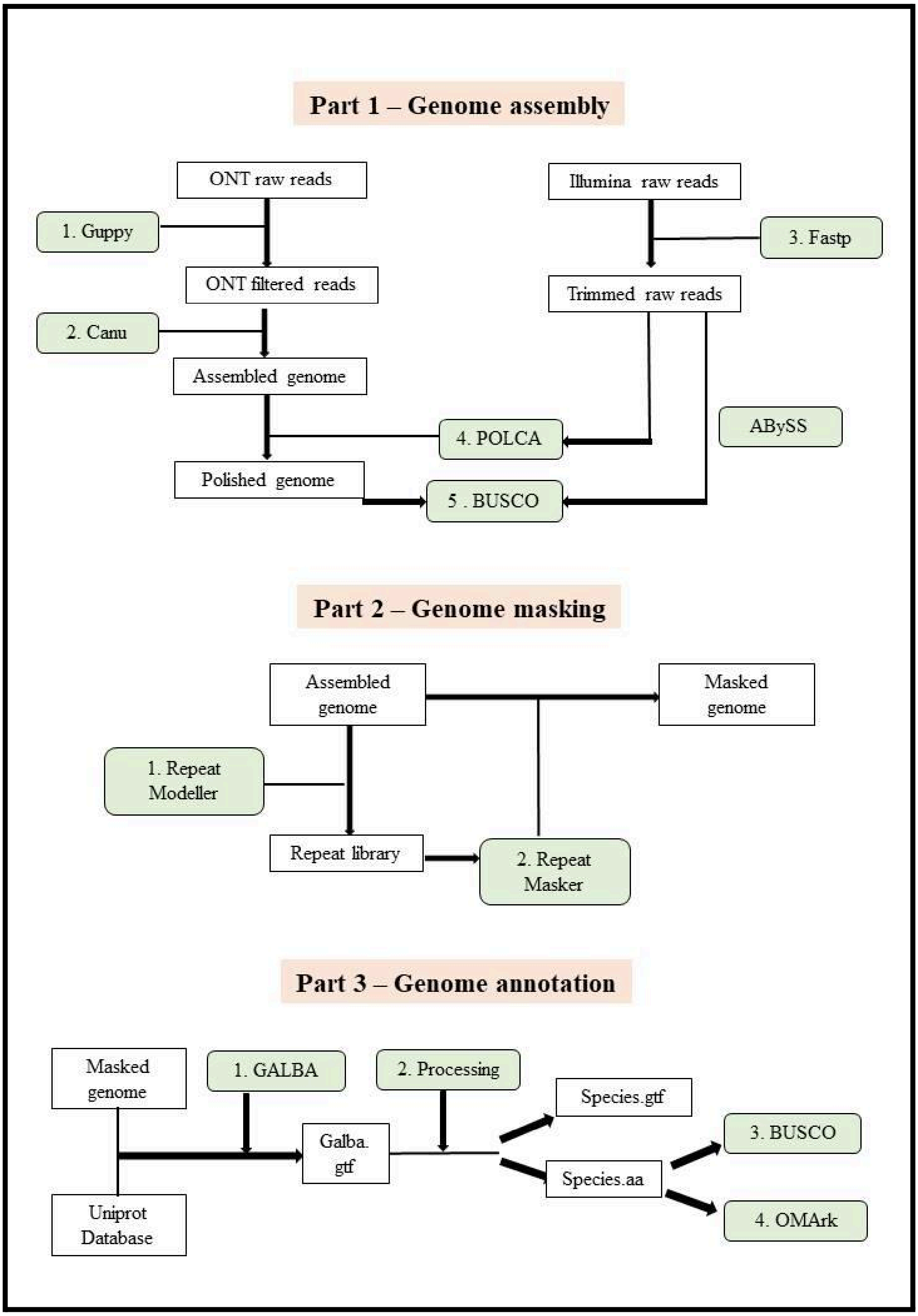

| Analyses | Tools and versions | Non-default parameters |
|---|---|---|
| Genome assembly using long-reads | Canu (v 2.2) | genomeSize=1g<br><br>rawErrorRate=0.30<br><br>correctedErrorRate=0.12 corMinCoverage=4<br><br>corMhapFilterThreshold=0.0000000002<br><br>corMhapOptions="--Threshold 0.80 --num-hashes 512 --num-min-matches 3 --ordered-sketch-size 1000 --ordered-kmer-size 14 --min-olap-length 2000 --repeat-idf-scale 50"<br><br>mhapMemory=192g<br><br>mhapBlockSize=500<br><br>ovlMerDistinct=0.975 |
| Genome assembly using short-reads | ABySS<br><br>(v 2.3.8) | k=96 B=25G j=48 |
| Correct short-reads | Fastp<br><br>(v 0.23.2) | --detect_adapter_for_pe<br><br>--trim_poly_g -5 -3<br><br>-l 51 |
| Polish long-read assembly with trimmed short-read data | Polca pipeline of Masurca<br><br>(v 4.1.0) | -t 50 -m 2G |
| Repeat masking | RepeatModeler (v 2.0.5) and | -BuildDatabase -name <path to polca corrected fa><br><br>-LTRStruct |

| | | |
|---|---|---|
| | RepeatMasker (v 4.1.5) | -nolow -s -e rmblast -xsmall -lib {custom library from RepeatModeler} |
| Gene prediction | GALBA (v 1.0.11) | -prot_seq=Aves_concat.fa |
| Genome annotation completeness analysis | BUSCO (v 5.7.1) | -lineage_dataset aves_odb10 |
| Genome annotation completeness analysis | OMArk (v 2.0.3) | |
| SNP and SVs estimation | MUmmer (v 3.23)<br><br>Assemblytics (v1.2.1) | nucmer -ref -query out.delta<br>show-snps out.delta -std.out<br><br>Assemblitics out.delta -prefix -max -min -block |
| Mitogenome assembly and annotation | GetOrganeller (v 1.7.7.1)<br>MITOS2 web server<br>Mitoz (v 3.6) | -genes GetOrganelle/LabelDatabase/animal_mt.fasta<br><br>mitoz all --data_size_for_mt_assembly 8,0 --clade Chordata --genetic_code 2 --species_name --fq1 --fq2 --skip_filter --requiring_taxa Chordata |
| Demographic history | PSMC (v. 0.6.5 | -N25 -t5 -r5 -p ‘25*2+4+4+6 |
| Quality check of the mapped reads | Qualimap (v.2.2.2) | bamqc |

**Supplementary Section 4: Supplementary results**

**Supplementary Table 1**: Details of genomic variation between Great Hornbill, Wreathed Hornbill, Rufous-necked Hornbill, and Malabar Pied Hornbill in comparison with published Great Hornbill genome (ASM2756380v1).

| Type of variation | Great Hornbill *Buceros bicornis* | Wreathed Hornbill *Rhyticeros undulatus* | Rufous-necked Hornbill *Aceros nipalensis* | Malabar Pied Hornbill *Anthracoceros coronatus* |
|---|---|---|---|---|
| Single nucleotide polymorphism (bp) | 1,925,217 | 53,080,775 | 52,485,788 | 63,684,412 |
| Insertions (50-500 bp) | 45737 | 955134 | 1098041 | 1024876 |
| Insertions(500-10000 bp) | 383674 | 4318782 | 2703125 | 2814869 |
| Deletions (50-500 bp) | 559466 | 2062227 | 2054137 | 1871811 |
| Deletions (500-10000 bp) | 558400 | 3847338 | 3886550 | 3991136 |
| Tandem expansion (50-500 bp) | 80570 | 132152 | 544376 | 249796 |
| Tandem expansion (500-10000 bp) | 401601 | 3627110 | 217329 | 288722 |
| Tandem contraction (50-500 bp) | 399357 | 197464 | 139128 | 183086 |
| Tandem contraction (500-10000 bp) | 285605 | 117489 | 68235 | 75973 |
| Repeat expansion (50-500 bp) | 951448 | 807778 | 549838 | 700392 |
| Repeat expansion (500-10000 bp) | 6466884 | 6435227 | 2699118 | 3345006 |

| | | | | |
|---|---|---|---|---|
| Repeat contraction (50-500 bp) | 1021228 | 1733464 | 1111422 | 1496590 |
| Repeat contraction (500-10000 bp) | 4391361 | 14168436 | 7850777 | 11121572 |

**Supplementary Table 2**: Mitogenome organization in Great Hornbill *Buceros bicornis* in comparison with published mitogenome

| Genes | names | from | to | strand | size_bp | size (Chen 2018) |
|---|---|---|---|---|---|---|
| tRNA-Phe | trnF(gaa) | 1 | 77 | 77 | + | 76 |
| 12S rRNA | rrnS | 76 | 1047 | 972 | + | 971 |
| tRNA-Val | trnV(tac) | 1046 | 1117 | 72 | + | 71 |
| 16S rRNA | rrnL | 1117 | 2712 | 1596 | + | 1580 |
| tRNA-Leu | trnL2(taa) | 2712 | 2786 | 75 | + | 74 |
| NADH1 | nad1 | 2797 | 3775 | 979 | + | 971 |
| tRNA-Ile | trnI(gat) | 3773 | 3845 | 73 | + | 72 |
| tRNA-Gln | trnQ(ttg) | 3857 | 3928 | 72 | - | 71 |
| tRNA-Met | trnM(cat) | 3927 | 3996 | 70 | + | 69 |
| NADH2 | nad2 | 3996 | 5037 | 1042 | + | 1042 |
| tRNA-Trp | trnW(tca) | 5036 | 5108 | 73 | + | 72 |
| tRNA-Ala | trnA(tgc) | 5109 | 5178 | 70 | - | 69 |
| tRNA-Asn | trnN(gtt) | 5181 | 5255 | 75 | - | 74 |
| tRNA-Cys | trnC(gca) | 5257 | 5324 | 68 | - | 67 |
| tRNA-Tyr | trnY(gta) | 5324 | 5393 | 70 | - | 70 |

| | | | | | | |
|---|---|---|---|---|---|---|
| COI | cox1 | 5394 | 6945 | 1552 | + | 1551 |
| tRNA-Ser | trnS2(tga) | 6936 | 7010 | 75 | - | 74 |
| tRNA-Asp | trnD(gtc) | 7012 | 7081 | 70 | + | 69 |
| COII | cox2 | 7082 | 7766 | 685 | + | 684 |
| tRNA-Lys | trnK(ttt) | 7767 | 7837 | 71 | + | 70 |
| ATP8 | atp8 | 7838 | 8006 | 169 | + | 168 |
| ATP6 | atp6 | 7996 | 8680 | 685 | + | 684 |
| COIII | cox3 | 8679 | 9463 | 785 | + | 784 |
| tRNA-Gly | trnG(tcc) | 9463 | 9532 | 70 | + | 69 |
| NADH3 | nad3 | 9532 | 9884 | 353 | + | 177 |
| tRNA-Arg | trnR(tcg) | 9886 | 9955 | 70 | + | 69 |
| NADH4L | nad4l | 9956 | 10253 | 298 | + | 297 |
| NADH4 | nad4 | 10246 | 11629 | 1384 | + | 1378 |
| tRNA-His | trnH(gtg) | 11624 | 11693 | 70 | + | 69 |
| tRNA-Ser | trnS1(gct) | 11693 | 11760 | 68 | + | 67 |
| tRNA-Leu | trnL1(tag) | 11760 | 11831 | 72 | + | 71 |
| NADH5 | nad5 | 11831 | 13646 | 1816 | + | 1836 |

| | | | | | | |
|---|---|---|---|---|---|---|
| CYTB | cob | 13654 | 14797 | 1144 | + | 720 |
| tRNA-Thr | trnT(tgt) | 14800 | 14870 | 71 | + | 70 |
| tRNA-Pro | trnP(tgg) | 14876 | 14947 | 72 | - | 71 |
| NADH6 | nad6 | 14955 | 15477 | 523 | - | 522 |
| tRNA-Glu | trnE(ttc) | 15481 | 15553 | 73 | - | 72 |
| D-loop | | | | | | 553<br>30 |

**Supplementary Table 3**: Mitogenome organisation of Wreathed Hornbill *Rhyticeros undulatus*

| Gene (Chen 2017) | name | from | to | size_bp | strand | size_bp (Chen2019) |
|---|---|---|---|---|---|---|
| tRNA–Phe | trnF | 111 | 183 | 73 | + | 72 |
| 12S rRNA | rrnL | 182 | 1156 | 975 | + | 1562 |
| tRNA–Val | trnV | 1155 | 1228 | 74 | + | 73 |
| 16S rRNA | rrnS | 1228 | 2827 | 1600 | + | 972 |
| tRNA–Leu | trnL2 | 2828 | 2902 | 75 | + | 74 |
| NADH1 | nad1 | 2915 | 3893 | 979 | + | 978 |
| tRNA–Ile | trnI | 3891 | 3964 | 74 | + | 73 |
| tRNA–Gln | trnQ | 3976 | 4047 | 72 | - | 71 |
| tRNA–Met | trnM | 4046 | 4115 | 70 | + | 69 |
| NADH2 | nad2 | 4115 | 5156 | 1042 | + | 1041 |
| tRNA–Trp | trnW | 5155 | 5230 | 76 | + | 75 |
| tRNA–Ala | trnA | 5241 | 5310 | 70 | - | 69 |
| tRNA–Asn | trnN | 5322 | 5397 | 76 | - | 75 |
| tRNA–Cys | trnC | 5405 | 5472 | 68 | - | 67 |

| | | | | | | |
|---|---|---|---|---|---|---|
| tRNA–Tyr | trnY | 5472 | 5543 | 72 | - | 71 |
| COI | cox1 | 5544 | 7095 | 1552 | + | 1551 |
| tRNA–Ser | trnS2 | 7086 | 7160 | 75 | - | 74 |
| tRNA–Asp | trnD | 7170 | 7239 | 70 | + | 67 |
| COII | cox2 | 7240 | 7924 | 685 | + | 679 |
| tRNA–Lys | trnK | 7925 | 7998 | 74 | + | 73 |
| ATP8 | atp8 | 7999 | 8164 | 166 | + | 165 |
| ATP6 | atp6 | 8154 | 8838 | 685 | + | 684 |
| COIII | cox3 | 8837 | 9621 | 785 | + | 784 |
| tRNA–Gly | trnG | 9621 | 9690 | 70 | + | 69 |
| NADH3 | nad3 | 9690 | 10042 | 353 | + | 352 |
| tRNA–Arg | trnR | 10044 | 10113 | 70 | + | 69 |
| NADH4L | nad4l | 10114 | 10411 | 298 | + | 297 |
| NADH4 | nad4 | 10404 | 11782 | 1379 | + | 1378 |
| tRNA–His | trnH | 11782 | 11851 | 70 | + | 69 |
| tRNA–Ser | trnS1 | 11851 | 11919 | 69 | + | 68 |
| tRNA–Leu | trnL1 | 11938 | 12009 | 72 | + | 71 |

| | | | | | | |
|---|---|---|---|---|---|---|
| NADH5 | nad5 | 12009 | 13830 | 1822 | + | 1821 |
| CYTB | cob | 13829 | 14972 | 1144 | + | 1143 |
| tRNA–Thr | trnT | 14975 | 15045 | 71 | + | 70 |
| tRNA–Pro | trnP | 15052 | 15123 | 72 | - | 71 |
| NADH6 | nad6 | 15130 | 15652 | 523 | - | 522 |
| tRNA–Glu | trnE | 15654 | 15727 | 74 | - | 73 |
| D-loop | OH | | | | | 2228 |

**Supplementary Table 4**: Mitogenome organization of Rufous-necked Hornbill *Aceros nipalensis*

| Gene | gene | from | to | size_bp | strand |
|---|---|---|---|---|---|
| tRNA–Glu | trnE(uuc) | 32 | 105 | 74 | + |
| NADH6 | ND6 | 106 | 628 | 523 | + |
| tRNA–Pro | trnP(ugg) | 636 | 707 | 72 | + |
| tRNA–Thr | trnT(ugu) | 713 | 783 | 71 | - |
| CYTB | CYTB | 785 | 1928 | 1144 | - |
| NADH5 | ND5 | 1927 | 3748 | 1822 | - |
| tRNA–Leu | trnL(uag) | 3748 | 3819 | 72 | - |

| | | | | | |
|---|---|---|---|---|---|
| tRNA–Ser | trnS(gcu) | 3818 | 3886 | 69 | - |
| tRNA–His | trnH(gug) | 3886 | 3955 | 70 | - |
| NADH4 | ND4 | 3950 | 5333 | 1384 | - |
| NADH4L | ND4L | 5326 | 5623 | 298 | - |
| tRNA–Arg | trnR(ucg) | 5624 | 5693 | 70 | - |
| NADH3 | ND3 | 5695 | 6047 | 353 | - |
| tRNA–Gly | trnG(ucc) | 6047 | 6116 | 70 | - |
| COIII | COX3 | 6116 | 6900 | 785 | - |
| ATP6 | ATP6 | 6899 | 7583 | 685 | - |
| ATP8 | ATP8 | 7573 | 7738 | 166 | - |
| tRNA–Lys | trnK(uuu) | 7739 | 7810 | 72 | - |
| COII | COX2 | 7811 | 8495 | 685 | - |
| tRNA–Asp | trnD(guc) | 8496 | 8565 | 70 | - |
| tRNA–Ser | trnS(uga) | 8568 | 8642 | 75 | + |
| COI | COX1 | 8633 | 10184 | 1552 | - |
| tRNA–Tyr | trnY(gua) | 10185 | 10254 | 70 | + |
| tRNA–Cys | trnC(gca) | 10254 | 10321 | 68 | + |

| | | | | | |
|---|---|---|---|---|---|
| tRNA–Asn | trnN(guu) | 10323 | 10398 | 76 | + |
| tRNA–Ala | trnA(ugc) | 10416 | 10485 | 70 | + |
| tRNA–Trp | trnW(uca) | 10497 | 10572 | 76 | - |
| NADH2 | ND2 | 10571 | 11612 | 1042 | - |
| tRNA–Met | trnM(cau) | 11612 | 11681 | 70 | - |
| tRNA–Gln | trnQ(uug) | 11680 | 11751 | 72 | + |
| tRNA–Ile | trnI(gau) | 11763 | 11836 | 74 | - |
| NADH1 | ND1 | 11834 | 12812 | 979 | - |
| tRNA–Leu | trnL(uaa) | 12825 | 12899 | 75 | - |
| 16S rRNA | l-rRNA | 12900 | 14504 | 1605 | - |
| tRNA–Val | trnV(uac) | 14504 | 14577 | 74 | - |
| 12S rRNA | s-rRNA | 14576 | 15552 | 977 | - |
| tRNA–Phe | trnF(gaa) | 15551 | 15622 | 72 | - |

**Supplementary Table 5**: Mitogenome organisation of Malabar Pied Hornbill *Anthracoceros coronatus*

| Genes | names | from | to | size_bp | strand |
|---|---|---|---|---|---|
| tRNA-Phe | trnF(gaa) | 1 | 76 | 76 | + |
| 12S | s-rRNA | 75 | 1050 | 976 | + |
| tRNA-Val | trnV(uac) | 1049 | 1122 | 74 | + |
| 16S | l-rRNA | 1122 | 2715 | 1594 | + |
| tRNA-Leu | trnL(uaa) | 2715 | 2789 | 75 | + |
| NADH | ND1 | 2798 | 3776 | 979 | + |
| tRNA-Ile | trnI(gau) | 3774 | 3847 | 74 | + |
| tRNA-Gln | trnQ(uug) | 3860 | 3931 | 72 | - |
| tRNA-Met | trnM(cau) | 3930 | 3999 | 70 | + |
| NADH | ND2 | 3999 | 5040 | 1042 | + |
| tRNA-Trp | trnW(uca) | 5039 | 5112 | 74 | + |
| tRNA-Ala | trnA(ugc) | 5125 | 5194 | 70 | - |
| tRNA-Asn | trnN(guu) | 5197 | 5272 | 76 | - |
| tRNA-Cys | trnC(gca) | 5274 | 5341 | 68 | - |
| tRNA-Tyr | trnY(gua) | 5341 | 5413 | 73 | - |
| cytochrome | COX1 | 5414 | 6965 | 1552 | + |
| tRNA-Ser | trnS(uga) | 6956 | 7030 | 75 | - |
| tRNA-Asp | trnD(guc) | 7032 | 7101 | 70 | + |
| cytochrome | COX2 | 7102 | 7786 | 685 | + |
| tRNA-Lys | trnK(uuu) | 7787 | 7858 | 72 | + |
| ATP | ATP8 | 7859 | 8027 | 169 | + |
| ATP | ATP6 | 8017 | 8701 | 685 | + |
| cytochrome | COX3 | 8700 | 9484 | 785 | + |
| tRNA-Gly | trnG(ucc) | 9484 | 9553 | 70 | + |

| | | | | | |
|---|---|---|---|---|---|
| NADH | ND3 | 9553 | 9905 | 353 | + |
| tRNA-Arg | trnR(ucg) | 9907 | 9976 | 70 | + |
| NADH | ND4L | 9977 | 10274 | 298 | + |
| NADH | ND4 | 10267 | 11650 | 1384 | + |
| tRNA-His | trnH(gug) | 11645 | 11714 | 70 | + |
| tRNA-Ser | trnS(gcu) | 11714 | 11782 | 69 | + |
| tRNA-Leu | trnL(uag) | 11781 | 11852 | 72 | + |
| NADH | ND5 | 11852 | 13698 | 1847 | + |
| cytochrome | CYTB | 13674 | 14817 | 1144 | + |
| tRNA-Thr | trnT(ugu) | 14820 | 14889 | 70 | + |
| tRNA-Pro | trnP(ugg) | 14895 | 14966 | 72 | - |
| NADH | ND6 | 14974 | 15496 | 523 | - |
| tRNA-Glu | trnE(uuc) | 15498 | 15570 | 73 | - |

**Supplementary Table 6**: OMark genome annotation statistics for Great Hornbill and Wreathed Hornbill

| OMArk statistics | Great Hornbill<br>*Buceros bicornis* | Wreathed Hornbill<br>*Rhyticeros undulatus* |
|---|---|---|
| **Completeness assessment (%)** | | |
| Single | 90.17 | 93.07 |
| Duplicated | 4.46 | 4.07 |
| Duplicated, Unexpected | 4.38 | 3.99 |
| Duplicated, Expected | 0.07 | 0.08 |
| Missing | 5.37 | 2.8 |
| **Consistency assessment (%)** | | |
| Total Consistent | 86.50 | 87.05 |
| Consistent, partial hits | 9.37 | 7.70 |
| Consistent, fragmented | 7.72 | 6.30 |
| Total Inconsistent | 4.09 | 4.25 |
| Inconsistent, partial hits | 0.92 | 0.79 |
| Inconsistent, fragmented | 1.61 | 1.47 |
| Total Contaminants | 0.00 | 0.00 |
| Total Unknown | 9.41 | 8.70 |
| **Species composition** | | |
| Clade | Neognathae (Strigiformes) | Neognathae |
| Number of associated query protein | 19310 (90.59%) | 19498 (91.30%) |

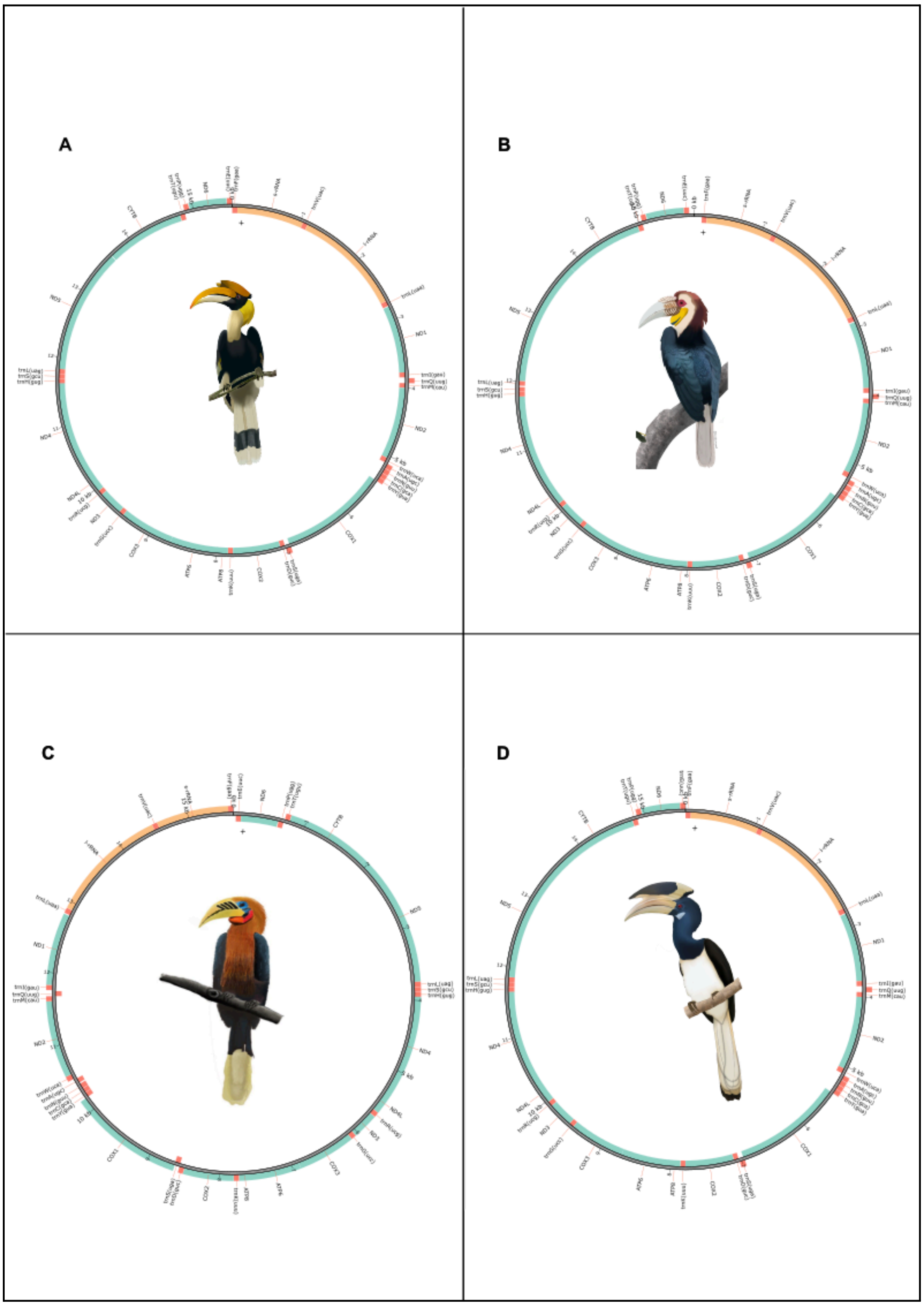


**Supplementary Figure 1**: Annotated mitochondrial genome plots for A. Great Hornbill, B. Wreathed Hornbill, C. Rufous-necked Hornbill, and D. Malabar Pied Hornbill. The genetic fragments in mitochondrial genome plots indicate the positions of protein-coding genes (green), rRNA (blue), and positions of tRNA (orange). (Illustrations by Bhagyashri Patwardhan)
